# MorphoGNN: Morphological Embedding for Single Neuron with Graph Neural Networks

**DOI:** 10.1101/2022.05.09.491257

**Authors:** Tianfang Zhu, Gang Yao, Dongli Hu, Chuangchuang Xie, Hui Gong, Anan Li

## Abstract

With the development of optical imaging systems, neuroscientists can now obtain large datasets of morphological structure at a single neuron scale positioned across the whole mouse brain. However, the enormous amount of morphological data challenges the classic approach of neuron classification, indexing and other analysis tasks. In this paper, we propose MorphoGNN, a single neuron morphological embedding based on the graph neural networks (GNN). This method learns the spatial structure information between the nodes of reconstructed neuron fibers by its nearest neighbors on each layer and captures the lower-dimensional representation of a single neuron through an end-to-end model. This model is composed of densely connected edge convolution (EdgeConv) layers and a double pooling operator, regularized with joint cross-entropy loss and triplet loss. An increasing population of the neighbor nodes meets the need of learning more information with features expanding at the deep layer. We tested the proposed embeddings on the neuron classification and retrieval tasks. Our method achieves competitive performance both on the general point cloud dataset and the neuron morphology dataset.

## Introduction

Rapid advances in the whole brain optical microscopy (1, 2) allow researchers to obtain extremely large amounts of brain slice images, where one mouse brain may generate several terabytes of data (3). Then the neuron fiber tracing methods (4–6) can reconstruct the whole structure of a single neuron fiber at a submicron resolution from the reconstructed 3D images. The reconstructed data usually includes a set of 3D points connected in the brain space (7), which describes the morphological structure of the neurons. These set of connected data points are converted to morphological features and the quantitative analysis of the neuron morphological features lays the foundation for neuron classifying by morphology and understanding their function (8–10).

To extract the morphological features, a series of neuronal morphometrics have been proposed, including the total neuron fiber length, the number of sections, the branch order, the radial distance, and other metrics (11). Classifiers can use these measurements to categorize neurons but the performance is limited due to the fact that a small difference in the local branch may change the neuron morphology substantially. Recently, researchers have focused on the learnable morphological features beyond these metrics. For example, the topological morphology descriptor (12) combines the topology of neuronal fibers with their spatial location, which allows it to distinguish neuron trees and different randomly generated tree structures. However, due to the intrinsic limitation of the morphometrics above, using them as the global feature is not appropriate. Cellular morphology neural networks (13) can also convert reconstructed neurons into vector representations. Specifically, the neuron is described by a series of 2D images from multiple view points, and then the representing features are extracted from the images by the convolutional neural networks. This method has been tested on electron microscopy data, but its effectiveness may be reduced when it is applied to the optical microscopic data due to the sparse subcellular structures.

We notice that the reconstructed neuron data presents a similar structure to the point cloud, where several processing and analysis methods (14, 15) have been proposed in recent years. Multi-view convolutional neural networks (16) demonstrates a collection of 2D views can accurately describe 3D objects. GNN (17, 18) captures the dependency relationship in the graph by information transfer between nodes, which is also introduced into the field of point cloud processing. (19) updates the graph structure between layers to learn the low-dimensional features of the point set, and then it can be applied to classification and segmentation tasks.

In this study, we propose MorphoGNN to learn the morphologic representation of a single neuron. The main contributions are shown as follows:

- We propose to use the GNN model to directly learn the morphologic information of a single neuron, where we can capture the low-dimensional global features of the neurons through a data-driven approach.
- This paper also introduces the K nearest neighbor (KNN) strategy to dynamically update the local graph, where the K value increases gradually with the increasing feature dimension. Dense connection, double pooling operator as well as joint loss function are introduced to the network architecture, which can improve the performance of the model effectively.
- We also provide a collection of neuron morphology datasets and compare our method with other baselines on the dataset in the neuron classification task, and neuron retrieval.

The basic principle, formula derivation, as well as network architecture of MorphoGNN are expounded in section Method. Section Experiment introduces the experiment datasets and preprocessing details, and presents the results of point cloud classification, neuron classification and neuron retrieval. Finally, the limitation of our method and the possible follow-up work are discussed in section Conclusion.

### Related work

#### Morphometrics

The development of large-scale and high-resolution micro-optical imaging technology (1–3) can obtain more complete morphological data of neuronal axons and dendrites, which give birth to various morphometrics describing neuronal topology. In addition to basic morphometrics such as the total neuron fiber length and the number of sections (11), recent morphological studies have focused on more fine and rich features. (20, 21) encodes the branches of axons and dendrites of nerve fibers, and calculates the similarity between trees by comparing the corresponding relationship of branch levels. (12) and (22) describe neuronal topology based on persistent homology and fiber evolution in brain space, respectively. In general, these morphometrics focus on a certain characteristic of neuron morphology, so researchers usually choose a combination of them to study a specific task.

#### Learning-based morphometrics

Learning-based methods have played a great role in biomedical fields, including brain region segmentation (23–25), medical image registration (26–29) and neuron reconstruction (30–32). However, these successful applications are based on regular data, such as medical images. Due to the irregularity of neurons, the existing deep learning technology can not directly act on morphological data, resulting in less work on morphological features based on learning method. Inspired by (16), (13) uses a set of images to describe a section of 3D nerve fibers, and then analyzes the image through a 2D convolution neural network to indirectly extract morphological features. This method tests a series of automatic tasks such as glia detection on electron microscope data. At present, for the automatic processing needs of massive reconstructed neurons, the morphological feature extraction method based on learning still needs more research.

## Method

We consider the reconstructed single neuron as an independent graph composed of a set of nodes. A standard neuron storage file format specifies the connections between nodes, ensuring that the entire neuron forms a tree structure. But the shape of each neuron is determined by nodes, regardless of these local connections. Hence each neuron morphologic structure can be viewed as a point cloud *P* = { *P*_*i*_ *R*^4^ = *x*_*i*_, *y*_*i*_, *z*_*i*_, *d*_*i*_} | *i* = 1, 2, …, *n*}, *x*_*i*_, *y*_*i*_, and *z*_*i*_ represent the 3D coordinates of these nodes, while *d*_*i*_ represents the diameter of the neuronal fiber at the node. Since the diameter measurements of neuronal fibers obtained by the current methods are related to the fluorescent strength and other factors, only spatial coordinates are considered in this study. The problem focused in this paper can be described in Eq. (1):

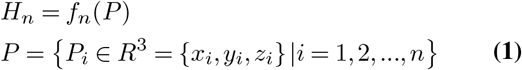

where *f*_*n*_ represent a embedding function. The function is supposed to capture the global features *H*_*n*_ of irregular neural fibers reconstructed:

Inspired by (19) and (14), we design the MorphoGNN that directly learns the representation embedding of the neuronal fibers, instead of manually designed morphometrics. To be specific, we update graph dynamically by the KNN method and learn node features for the higher dimensions continuously, then use the double pooling operator to capture the global features of neuronal fibers. In the process, the amount of information of network learning keeps increasing to cope with the increasing node characteristic dimension. The following are detailed explanations of the key elements of the proposed method.

### Local Graph Update

The points in the point cloud form a graph with the nearest K points, then new features of central node are learned from its neighbor nodes. As node features are updated, the graph is dynamically recomputed between layers. A local graph *G*^*l*^ at layer *l* can be defined as in Eq. (2):

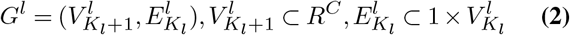

where *C* and *K* represent the dimension of node feature and the number of neighbors are selected at layer *l* respectively. As shown in Fig. 1, we set the K value to increase as the layer deepens so the local graph is also growing in size, which can be calculated by Eq. (3):

**Fig. 1.**
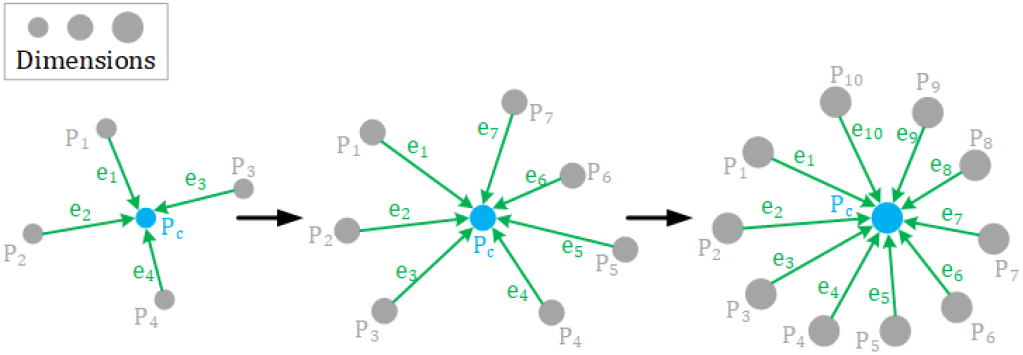
K value increases with node feature expanding. Diameters of points represent their feature dimensions. As node features expand, each node needs to learn from more neighbors.The dimension of node features in our network are {32,64,128,256} with K value {8,16,32,64}.

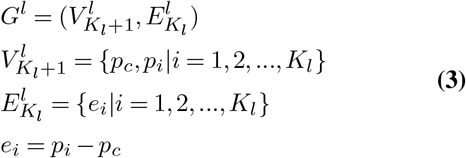
where *e*_*i*_ is the directed edge from one of the neighbor nodes *p*_*i*_ to the updating node *p*_*c*_.

### Local Feature Update

The initial feature of each node for a single neuron is a 3D (*x*_*i*_, *y*_*i*_, *z*_*i*_) vector, and the local features are updated to a higher dimension via the EdgeConv (19). For a local graph, the local feature of the updating point *p*_*c*_ can be generated by Eq. (4):

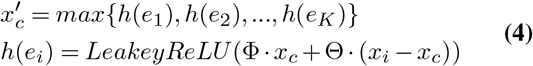
where Φ and Θ are trainable matrices, while *h*(*e*_*i*_) represents the hidden vector of the edge *e*_*i*_ from the neighbor node *p*_*i*_ to *p*_*c*_. The new local feature 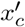 can be obtained through a max-pooling operation for all edge vectors. Since *e*_*i*_ is combined of the old local feature *x*_*c*_, initially coordinates, and the morphologic information of the neighborhood 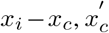 can be considered as the morphological features of neurons in another representation.

### Morphological Embedding by Pooling

The dimension *C* increases rapidly after a few iterations of updating on the node features *H*_*p*_. To improve the local feature utilization, the global feature of neurons is computed by two kinds of pooling operators for node features, which could be described as shown in Eq. (5):

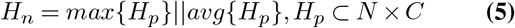
where *H*_*n*_ and *N* denotes the features of the single neuron and the number of points respectively, while ‖ represents the concatenation. Max-pooling captures the prominent feature, while average-pooling preserves the overall feature strength. Their outputs are concatenated so the final dimension of *H*_*n*_ is 2 × *C*.

### Classification and Indexing of Neurons

In this paper we use two tasks with the neuron features to demonstrate the proposed embedding method: neuron classification and its retrieval. For classification, we aim to find a function *f*_*c*_ that maps the features into a probability distribution *P*_*d*_ as defined in Eq. (6):

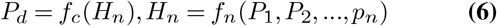
For the neuron retrieval, the actual task is to calculate the similarity *S* of neuron features in pairs, and retrieve the related neurons and rank the candidates from high similarity to low similarity. The neuron similarity function can be defined as in Eq. (7):

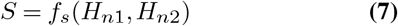
where *f*_*s*_ could be the cosine similarity or Minkowski distance.

### Joint loss function

In addition to the cross-entropy function commonly applied in classification tasks, we also use triplet loss (33) to deal with complex and diverse neurons. Considering the morphological embedding *h*_*a*_ of a single neuron, *h*_*p*_ and *h*_*n*_ are defined to represent the same and different embeddings in the training batch respectively. Triplet loss *L*_*t*_ can be calculated by Eq. (8):

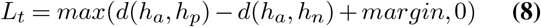

where *d* is a function to calculate the distance between two embeddings, usually euclidean distance. And *margin* is a constant greater than 0, making *d*(*h*_*a*_, *h*_*p*_) smaller and *d*(*h*_*a*_, *h*_*n*_) larger. As described in Eq. (8), triplet loss shortens the distance of similar morphological embeddings and lengthens the distance of heterogeneous morphological embedding, so as to identify neurons with small differences. The total criterion *L* can be expressed as the weighted sum of two loss functions:

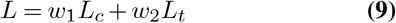
where *L*_*c*_ represent the cross-entropy loss function, *w*_1_ and *w*_2_ are the weights of the two loss functions.

### Network Architecture

Our network architecture is an improved version of DGCNN (19). In general, the dimension of neuron nodes features is increased from 3 to 256 by four EdgeConv layers, and then aggregated into the global features of nerve fibers through a pooling module. Finally, the classification results are output by two fully connected layers(See Fig. 2).

**Fig. 2.**
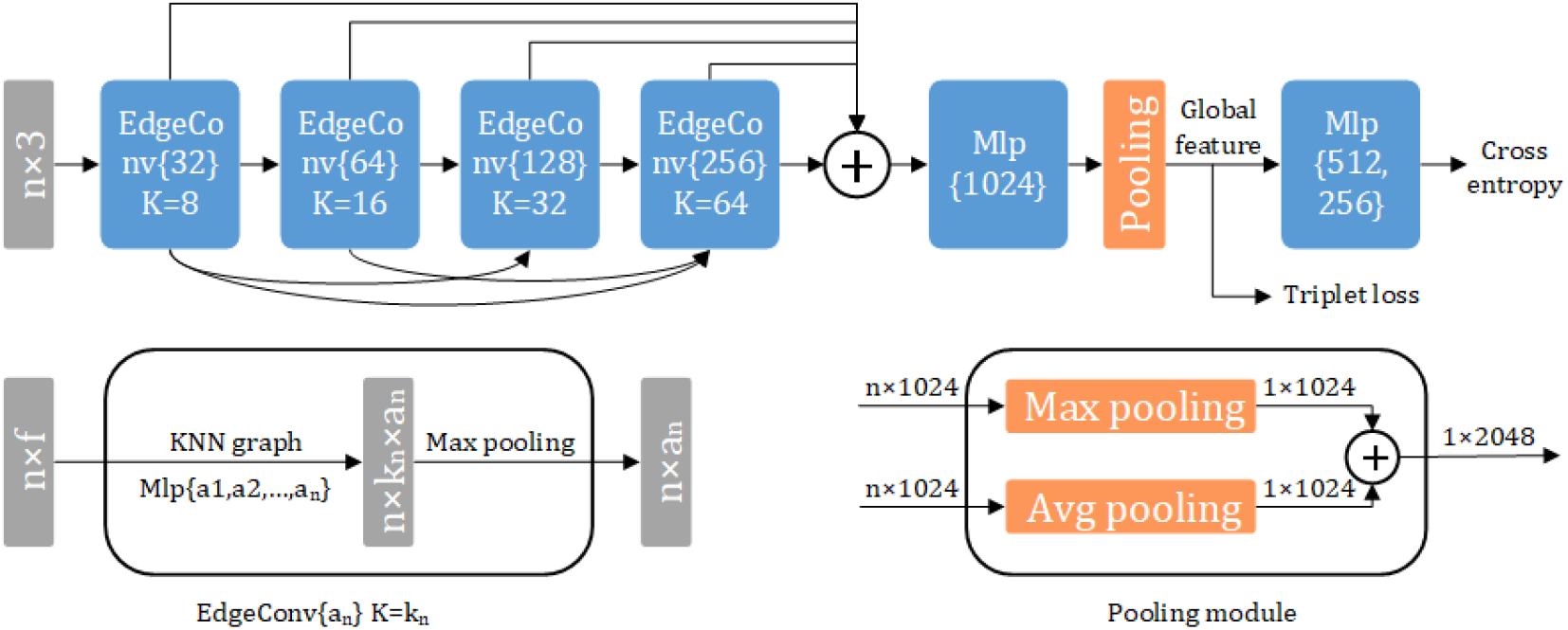
Network architecture. Our model takes the 3D coordinates of *n* points as input and outputs the classification scores of neurons. EdgeConv{*N*} denotes that after this EdgeConv layer, node features are expanded to *N* dimension and the value of *K* indicates how many neighbors are required per node in this layer. Mlp represents for multi-layer perceptron while ⊕ means concatenation operation. Morphological embedding is combined of the outputs of max-pooling and average-pooling layer.

A significant difference is that we adopt an increasing K value, while the K value of DGCNN is a global hyperparameter. More topological information is needed With the increasing node feature dimension. Therefore, each node needs to aggregate information from a wider range of neighbors to update its own characteristics. Furthermore, we add a structure that is similar to a dense connection. The input of each EdgeConv layer comes from the output of each previous EdgeConv. The output of the last EdgeConv layer is pooled and aggregated into the global feature vector. In order to get abundant global features, the output of the last EdgeConv layer goes through a maximum pooling layer and an average pooling layer respectively, then their outputs are concatenated into the final feature vector of nerve fibers.

General point cloud data, such as the ModelNet10/40 dataset (34), could be classified with a good performance by the softmax function output and the cross-entropy loss. However, there is no significant difference between the neuron of different types. Even the same neuron will show morphological diversity due to the difference in reconstruction methods, imaging resolution, as well as parameter settings. Thus triplet loss (33) and cross-entropy are used jointly to provide a stronger constraint on the network output.

#### Experiment

In this experiment section, we describe the data and training procedure for the proposed MorphoGNN and also show the performance of the model along with other baseline models.

### Data Description

We compare the network performance on point cloud dataset, and also on the collected neuron morphology dataset. The point cloud dataset is ModelNet40 (34), which includes 40 types of common objects, such as cars, airplanes, and guitars. Every single model in ModelNet40 is composed of 2048 points, where the 3D coordinates as the initial feature.

Besides the reconstructed 1393 neurons from 7-class on NeuroMorpho.org (35) are collected for our experiment and we denote the dataset as Neuro7. The dataset contains morphological data from several groups, of wild-type mouse brains, and using nonfluorescent stainings. However, the raw morphological data is not standardized due to the different sources of the contributors. As shown in Table. 1, we selected seven representative types of mouse neuron cells, each type of which is the result of contributions from multiple research groups. Aspiny neurons come from three groups (36–38). Spiny neurons are all from (36). (39) contributes all of the amacrine and bipolar neurons. Stellate neurons also consist of three groups (40–42), while basket neurons are from (43) and (44). Finally, pyramidal neurons are totally taken from four groups (45–48). The number of points that make up a neuron varies from 1 × 10^3^ to 6 ×10^3^. These two datasets are divided into a verification set and training set according to the ratio of 3 : 7.

**Table 1.** 7-class neuronal morphological data

| Neuron class | Amacrine | Aspiny | Basket | Bipolar | Pyramidal | Spiny | Stellate |
| --- | --- | --- | --- | --- | --- | --- | --- |
| Sample size | 232 | 98 | 130 | 355 | 239 | 247 | 92 |
| The min number of points | 1617 | 1118 | 1072 | 1163 | 1139 | 1468 | 1099 |
| The max number of points | 5719 | 5986 | 5354 | 5462 | 5701 | 5690 | 4603 |
| The mean number of points | 3614 | 2873 | 2065 | 2117 | 2002 | 2920 | 1953 |

### Morphological Data Preprocessing

The padding strategy eliminates the adverse effect caused by the inconsistent number of points per sample and makes it possible to group different neurons into a mini-batch to accelerate training. We use a fixed number of points (6 ×10^3^ in this work), which is greater than the maximum number of points, as the padded number of points per sample. The list of points for each neuron is appended with the triplet (0, 0, 0) until the size of the list is 6 × 10^3^. Although the padding changes the distribution of the points for each neuron, subsequent experiments show that it does not significantly affect network performance.

Normalization is another necessary step in preprocessing the neuronal morphological data. The absolute location of neurons in brain space leads to a wide difference between the coordinates of the reconstructed points. The following formula Eq. (10) is used to normalize the neuron point coordinates:

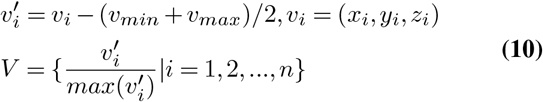

where *v*_*max*_ and *v*_*min*_ are the maximum and minimum of initial coordinates *v*_*i*_ respectively, while *V* represents the normalized coordinates set. We firstly find the smallest unit cube core that could surround the original neuron, take it as the new origin, and then normalize the new coordinate. Fig. 3 shows that the normalized neuron retains its original shape, with its center as the spatial origin, and the coordinate value of each point is in (−1, 1).

**Fig. 3.**
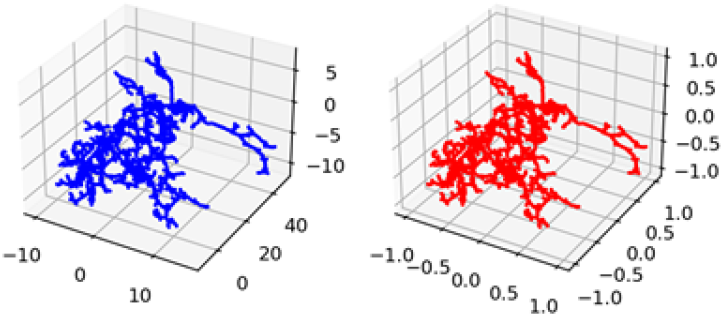
Normalization for neurons. **Blue** is the original neuron; **Red** is the normalized neuron. Raw coordinates are experiment dependent. After the normalization, they are scaled to range (−1, 1), with the origin located in the geometric center of the neuron.

### Implementation Details

Here we explain the training process of our network. We use the Adam optimizer (49) with a 10^−3^ learning rate and 0.9 momentum to train the whole network. Every 20 epochs, the learning rate drops by half. For ModelNet40 dataset, our network only uses the same crossentropy loss function as other baseline models. However, for morphological data, cross-entropy and triplet loss are used jointly, with a weight ratio of 1 : 1. The morphological embedding dimension learned by other methods is 1024, while our networks is 2048 due to the double pooling module. In addition, all of our experiments are run on one NVIDIA RTX 5000 GPU with 16GB of graphic memory.

### Classification on Morphological Dataset

We follow the settings in the ModelNet40 experiment and evaluate the effects of several methods on the morphological data for neuron classification. Here our network uses cross-entropy and triplet loss (triplet) for joint training, other methods remain the same. We also present the performance of our method using cross-entropy loss alone (baseline). Neuron classification based on the traditional morphometrics are also included for comparison. Five basic neuronal morphological parameters, including number of branches, number of leaf nodes, maximum branch orders, maximum radial distance, and total length form the feature vector of each neuron. These measurements are then used for neuron classification through a simple multi-layer perceptron network with the same parameters as the features extracted by other methods.

All the processing methods for point cloud models are less effective on morphological dataset, shown in Table 2. Our network still provides the best performance with the highest overall accuracy(85.58%), about 6.49% ahead of Point-Net++. The mean class accuracy of each method is significantly lower than the overall accuracy due to the difference between classes is not as obvious as that of point cloud and the class population is not balanced well. However, our method reaches a competitive 79.45% mean class accuracy, which improves around 5.3% over the model with cross-entropy loss, suggesting that joint loss plays an important role in a better performance. In addition, the discrimination power of morphometrics is also good, which achieves a higher overall accuracy(72.98%) than Pointnet.

**Table 2.** Classification on Neuron7

| Method | MA(%) | OA(%) |
| --- | --- | --- |
| PointNet | 58.65 | 43.57 |
| PointNet++ | 71.17 | 79.09 |
| DGCNN(k=20) | 70.61 | 79.33 |
| DGCNN(k=30) | 65.19 | 76.20 |
| DGCNN(k=64) | 67.71 | 76.20 |
| Morphometrics | 52.14 | 72.98 |
| Ours(baseline) | 74.14 | 81.49 |
| Ours(triplet) | 79.45 | 85.58 |

T-distributed stochastic neighbor embedding (T-SNE) is used to visually display the features extracted by these methods(Fig. 4). A single point in the figures represents the feature of a single neuron after dimensionality reduction. It is obvious that the features extracted by our method are more discriminative than those extracted by other methods. Features of the same class extracted by our method are close to each other and far away from other classes. However, some morphologically similar categories are not clearly distinguished, such as spiny, aspiny, basket, and stellate. It is worth noting that neurons described by five morphometrics clustered in a small area of the figure because of its low-dimension but still shows some discrimination.

**Fig. 4.**
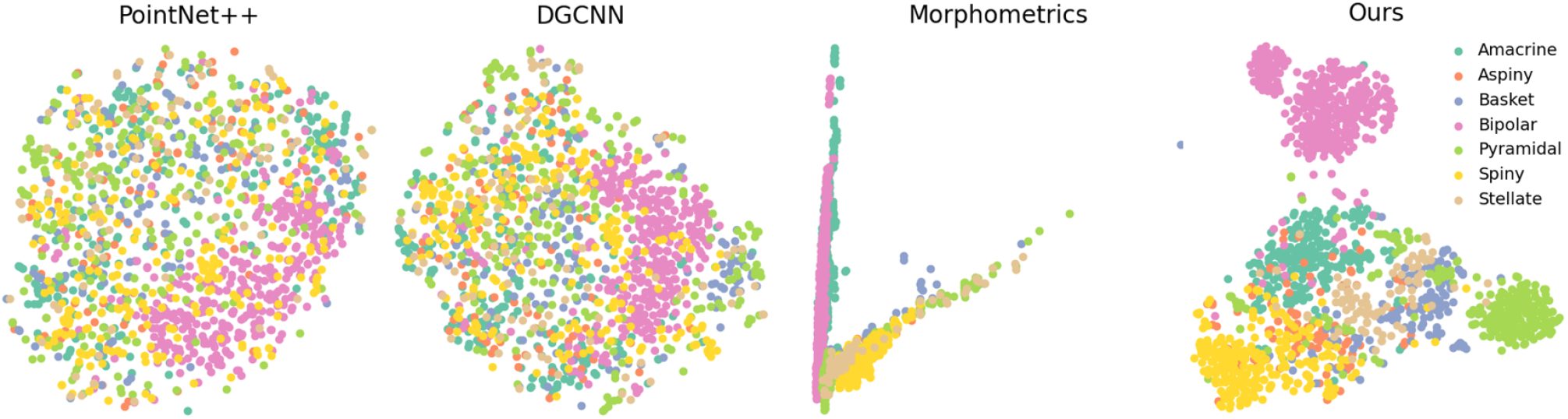
T-SNE visualization of different features. The features of each neuron are reduced to 2D and shown as points in the figure. Colors represent the corresponding categories of the neurons.

### Retrieval on Morphological Dataset

A morphological neuron database requires to support neuron indexing and retrieving, which can be based on the embedding similarity between neurons. Morphological data are divided into two versions according to whether the padding strategy is adopted or not. Then different methods are used to extract the features one by one to build the database. Table. 3 shows the time cost and retrieval accuracy required to build the database of different features. The accuracy of each retrieval operation is defined as the proportion of the same species among the ten most similar neurons retrieved, which is calculated according to the cosine similarity. According to Table 3, the padding strategy has little impact on the model performance. Our method achieves the highest overall accuracy(77.32%) and means class accuracy(67.82%) respectively. The time it takes to build the database is less than PointNet++ and more than DGCNN.

**Table 3.** Retrieval on Morphological Dataset

| Method | Embedding dimension | Padding |  |  | No Padding |  |  |
| --- | --- | --- | --- | --- | --- | --- | --- |
|  |  | Time(s) | MA(%) | OA(%) | Time(s) | MA(%) | OA(%) |
| PointNet | 1024 | 26.77 | 29.20 | 38.49 | 20.32 | 27.41 | 36.05 |
| PointNet++ | 1024 | 849.9 | 29.55 | 38.56 | 500.3 | 29.05 | 38.25 |
| DGCNN | 1024 | 112.4 | 30.67 | 39.94 | 47.07 | 31.52 | 41.26 |
| Morphometrics | 5 | - | - | - | 40.06 | 48.89 | 60.85 |
| MorphoGNN | 2048 | 231.1 | 67.82 | 77.32 | 93.65 | 66.29 | 76.23 |

Fig. 5 shows the five most similar neuron fibers retrieved from the same neuron fiber by the five methods. Due to the individual differences and the insignificant inter-class differences, the retrieval results of other methods are mixed with various kinds of neurons. But our method retrieves four neurons of the same class with one from the other class.

**Fig. 5.**
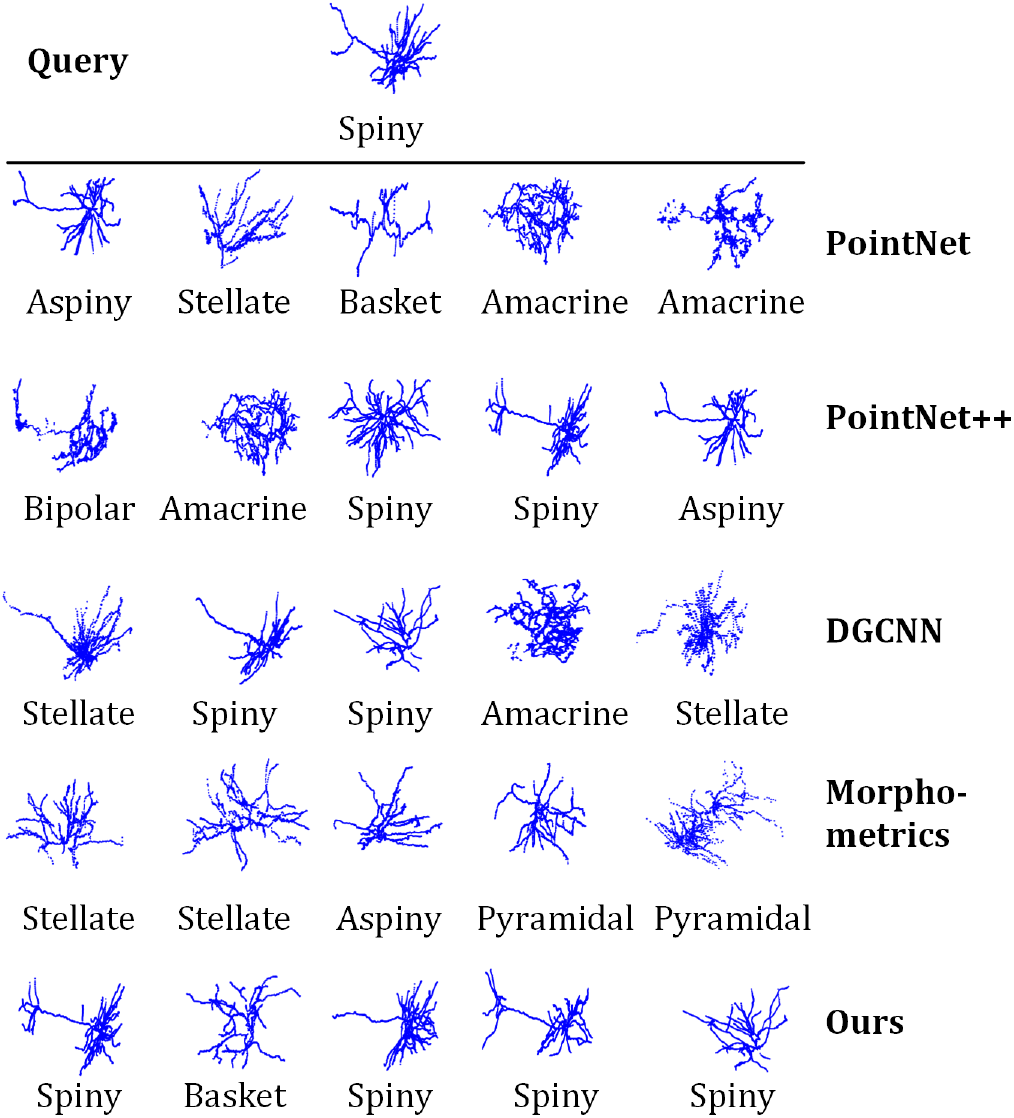
Retrieval results of five methods. **Left** is the randomly selected query neuron and **right** are the five most similar neurons retrieved, while texts below are the category to which these neurons actually belong.

### Robustness test

Due to some unavoidable defects in the process of imaging and reconstruction, some incomplete and inaccurate morphological data are often obtained. The cost of discarding all these samples is unbearable, so the algorithms are expected to be robust to low-quality neurons. In this section, we discuss the performance of MorphoGNN against two attacks, including point dropping and Gaussian noise. The former randomly drops some points in a single neuron to simulate the incomplete reconstruction, while the latter adds point-wise noise to make the reconstructed neuron shape deviate from the reality. We attacked the neurons in the testset to varying degrees(Fig 6), and then recorded the classification accuracy of the pretrainied MorphoGNN.

**Fig. 6.**
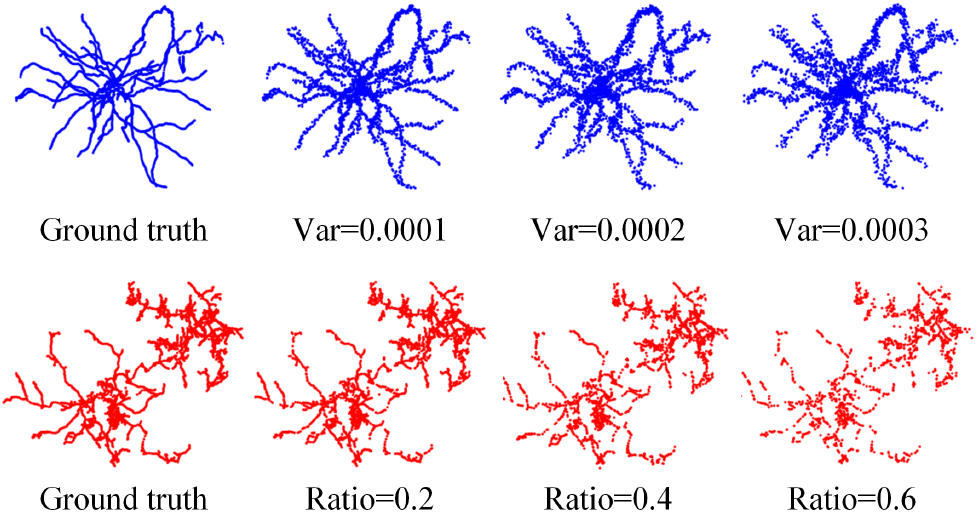
Neurons with point dropping and Gaussian noise. The **up** is Gaussian noise with different variance, and the **bottom** is point dropping with different ratio.

Fig 7 shows the overall accuracy and mean-class accuracy of MorphoGNN in the face of different degrees of point dropping and Gaussian noise attacks. It can be clearly seen that the overall accuracy and mean-class accuracy illustrate the same downward trend, indicating that the network performance is generally weakened when the morphology of neurons is destroyed. However, the performance degradation of MorphoGNN in the face of point dropping is slower than that in the face of random noise. When the variance of Gaussian noise increases to 1.5 × 10^−4^, the overall accuracy and average accuracy drop to less than 80% of the initial network, which is equivalent to discarding 35% of the points. The better robustness of MorphoGNN to point discarding may be due to the fact that many zero points are filled in during data preprocessing, which makes the network less sensitive to point dropping than random noise. Overall, experimental results show that low-quality neurons will significantly weaken the network performance, thus improving the network robustness is necessary to study in the future.

**Fig. 7.**
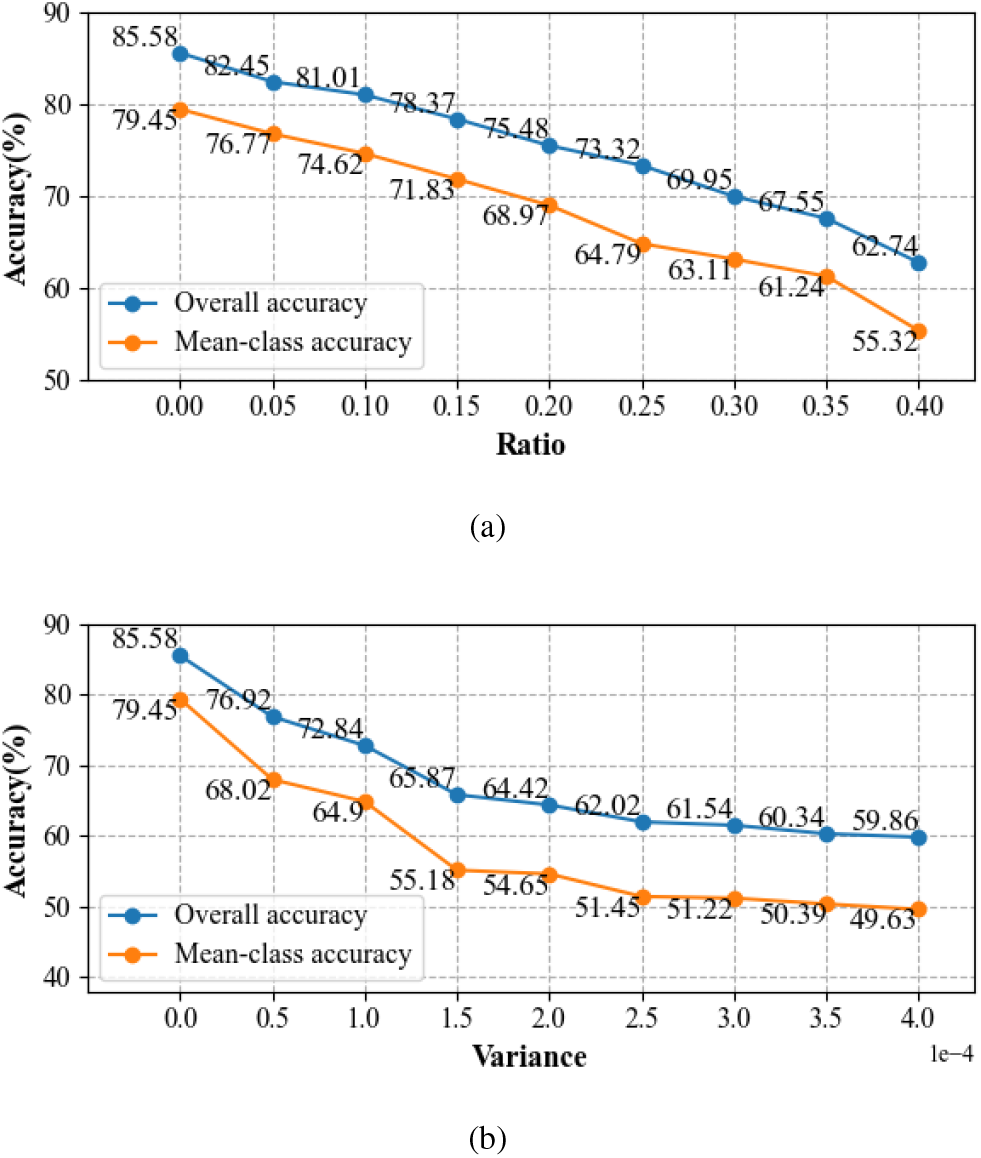
Influence of two attacks on MorphoGNN. The abscissa of **(a)** is the ratio of discarded points, while that of **(b)** is the variance of Gaussian noise.

## Conclusion

We introduce MorphoGNN, a method to obtain the morphological embedding of the reconstructed neuron fibers based on a proposed GNN, in order to meet the need for automatic analysis of massive morphology data. We show that the augmented local graphs with increasing node features can enhance the network performance, not only for neuronal data, but also for general point clouds. The performance of our network is tested on general point cloud models and we demonstrate the effectiveness of neuron classifying, and retrieving on a morphology dataset containing 1393 neurons.

### Limitations and future work

We treat the neuron fibers as point clouds, which provides convenience for applying GNN, but ignores the specific differences among neuron subcellular structures such as dendrites, axons, and cell bodies. In addition, this paper only focuses on the single neuron, while neural circuits composed of multiple neurons are the basis of brain function. Learning the connections between neurons will contribute to future understanding of brain function and stimulate the development of the brain-inspired intelligence.

## Acknowledgements

We thank Yue Guan from Huazhong University of science and technology for the helpful comments on experiments. This work was financially supported by the National Science and Technology Innovation 2030 Grant (2021ZD0201002) and the National Natural Science Foundation of China (T2122015).

